# A multi-omics atlas of the human retina at single-cell resolution

**DOI:** 10.1101/2022.11.21.517389

**Authors:** Qingnan Liang, Xuesen Cheng, Jun Wang, Leah Owen, Akbar Shakoor, John L. Lillvis, Charles Zhang, Michael Farkas, Ivana K. Kim, Yumei Li, Margaret DeAngelis, Rui Chen

## Abstract

Cell classes in the human retina are highly heterogeneous with their abundance varying by several orders of magnitude. Although previous studies reported the profiles of the retinal cell types as the transcriptome level, there is no study regarding the open-chromatin profiles at a similar resolution. Here, we generated and integrated a multi-omics single-cell atlas of the adult human retina, including over 250K nuclei for single-nuclei RNA-seq and 137K nuclei for single-nuclei ATAC-seq. Through enrichment of rare cell types, this single cell multiome atlas is more comprehensive than previous human retina studies. Cross species comparison of the retina atlas among human, monkey, mice, and chicken revealed relatively conserved and non-conserved types. Interestingly, the overall cell heterogeneity in primate retina decreases compared to that of rodent and chicken retina. Furthermore, integrative analysis of the single cell multi-omics data identified 35k distal cis-element-gene pairs with most of these cis-elements being cell type specific. We also showed that the cis-element-gene relationship in different cell types within the same class could be highly heterogenous. Moreover, we constructed transcription factor (TF)-target regulons for over 200 TFs, partitioned the TFs into distinct co-active modules, and annotated each module based on their cell-type specificity. Taken together, we present the most comprehensive single-cell multi-omics atlas of the human retina as a valuable resource that enables systematic in-depth molecular characterization at individual cell type resolution.

## Introduction

The vertebrate retina is a multi-layer neuronal structure that converts light to electrical signals that are transmitted to the brain^1^. The retina is composed of six major neuronal classes, rod, cone, bipolar, horizontal, amacrine, and retinal ganglion cells (RGCs), along with several non-neuronal cell types, such as Müller glia^1,2^. Although the overall structure and major cell classes of the retina are similar across all vertebrates, there are significant differences in cytoarchitecture, synaptic connections, and the retinal vascular bed between species as well as a complete absence of the macula in most non-primate species^3,4^. Recent single-cell transcriptomic studies of the retina have consistently indicated that the number and classification of amacrine and RGC types vary greatly across different species, providing a potential underlying mechanism of the retinal structure, circuitry, and functional divergence across species^5–8^. The human retina is comprised of a highly heterogeneous population of cells and it contains an estimated 60 neuronal types based on morphology, function, and most recently, single-cell transcriptional profiles^5,9–11^. The abundance of each cell type is highly variable and ranges from over 60% to less than 0.05% of the cell population. As a result, a combination of both an increase in the number of cells profiled and targeted enrichment of rare cell types is required to build a comprehensive cell atlas of the human retina.

Beyond single-cell transcriptomics, obtaining the epigenomic landscape at single-cell resolution is critical for gaining insights into the dynamics of gene expression regulation and identifying candidate regulatory elements and variants that affect transcription. Assay for Transposase-Accessible Chromatin sequencing (ATAC-seq) has emerged as an ideal method for profiling open chromatin regions, which contain most of the active regulatory elements for gene expression, in cells^12^. Recent advancements have made it possible to perform ATAC-seq to profile open chromatin regions for individual cells^13,14^. By combining single-nuclei ATAC-seq data with single-nucleus transcriptomics data, it is possible to systematically map gene cis-regulatory elements (CREs) for each cell type and elucidate the gene-regulatory landscape of the retina, which will be critical for identifying and prioritizing non-coding mutations in CREs associated with retinal disorders^15,16^. Furthermore, a large portion of GWAS hits have mapped to potential gene regulatory regions rather than the coding region of the genome^17,18^. Thus, obtaining a high-quality chromatin landscape along with transcriptomic profiles at single-cell resolution is a critical step toward better characterization of human retina biology and diseases at the molecular level. Several recent approaches reported single-cell level open chromatin profiles with human retina^19–22^, however none of these studies included extensive cell type classification, so in general, the open chromatin profiling for retina is still at the cell class level, instead of cell type level.

In this report, we built a single-cell multi-omics map of the human retina by profiling transcriptomes of over 250,000 nuclei and open chromatin from over 137,000 nuclei. In total, over 70 cell types, including 68 neuronal types, were identified, representing the most comprehensive map of the human retina to date. By integrating the single-nuclei ATAC-seq (snATAC-seq) and single-nuclei RNA-seq (snRNA-seq) data, open chromatin profiles were identified for each cell type. Cross-species comparison between humans, monkeys, mice, and chicken revealed various levels of similarity for the different cell types among these species. Counter-intuitively, we found that the sequence conservation of differential accessible regions could not explain the relative conservation of the cell types. Candidate CREs were identified and most of them showed strong cell type specificity. We also showed that the regulatory relationship between genome regions and genes are only partially shared among cell types, which suggested that evaluation of gene regulation at type level is required. Lastly, we constructed TF-target regulons and demonstrated that the TFs could potentially work in modules. We were able to annotate the TF modules and infer the hub regulators for the modules. In summary, our study has generated the most comprehensive single-cell multi-omics atlas that enables the in-depth characterization of the human retina at specific cell type resolution.

## Results

### The single-cell multiomics atlas of human retina

To generate a comprehensive cell atlas and identify most cell types of the human retina, we performed snRNA-seq and snATAC-seq on six and twenty-three healthy donors, respectively (Table 1-2). It is known that over 60 cell types exist in the primate retina^5^. Since the distribution of retinal types is not uniform, some of the cell types are exceedingly rare. For instance, at least 18 RGC types have been classified in the human retina based on morphology^23^, but the total number of RGCs only account for about 1% of the cell population in the retina^24^. In addition, the proportions of RGC types differ significantly, and some RGC types, such as intrinsically photosensitive RGCs (ipRGCs), make up less than 0.01% of retinal cells^25^. Similarly, it has been estimated that over 30 types of Amacrine cells (ACs) exist in the primate retina^5^. Therefore, to profile these rare cell types, to enrich RGCs and ACs before profiling could be very necessary.

**Table 1.** Information of the donors from whom the retina tissue was obtained and profiled.

| Donor-ID | Ethnicity | Sex | Age | Total Cell Number | Central | Peripheral | Peripheral (sorted) |
| --- | --- | --- | --- | --- | --- | --- | --- |
| D001-12 | European | F | 78 | 17664 | 0 | 0 | 17664 |
| 19D013 | European | F | 78 | 45905 | 41345 | 0 | 4560 |
| 19D014 | European | M | 84 | 45118 | 38497 | 0 | 6621 |
| 19D015 | Hispanic | M | 73 | 101089 | 57579 | 8283 | 35227 |
| 19D016 | European | M | 83 | 19645 | 19645 | 0 | 0 |
| 17D013 | European | M | 65 | 5470 | 0 | 0 | 5470 |

**Table 2.** Donor information and number of nuclei sequenced from each sample for single-nuclei ATAC-seq.

| Donor-ID | Ethnicity | Sex | Age | Total Cell Number |
| --- | --- | --- | --- | --- |
| 19D013 | European | F | 78 | 21085 |
| 19D014 | European | M | 84 | 7334 |
| 19D016 | European | M | 83 | 16255 |
| 19_D007 | Caucasian | M | 66 | 3577 |
| 19_D003 | Caucasian | M | 86 | 2606 |
| 19_D005 | Hispanic | M | 53 | 6527 |
| 19_D006 | Caucasian | M | 90 | 5526 |
| 19_D008 | Hispanic | F | 64 | 4489 |
| 19_D009 | Hispanic | M | 68 | 4815 |
| 19_D010 | Caucasian | F | 88 | 5334 |
| 19_D011 | Caucasian | F | 90 | 7137 |
| 19_D019 | Caucasian | F | 82 | 5570 |
| D005_13 | Caucasian | M | 80 | 4266 |
| D009_13 | Asian | M | 85 | 3854 |
| D013_13 | Caucasian | M | 69 | 4480 |
| D017_13 | Caucasian | M | 65 | 5295 |
| D018_13 | Caucasian | F | 91 | 1693 |
| D019_13 | Caucasian | M | 82 | 6281 |
| D021_13 | Caucasian | M | 86 | 2408 |
| D026_13 | Caucasian | M | 81 | 6632 |
| D027_13 | Caucasian | M | 77 | 4606 |
| D028_13 | Caucasian | M | 67 | 8018 |

NeuN is a nuclear envelope protein encoded by the *RBFOX3* gene that has been reported to be highly expressed in a subset of neuronal cells^26^. In the mouse retina, NeuN was shown to be highly expressed in RGCs and ACs but lowly expressed in photoreceptor^27^. Antibody staining showed that NeuN showed a similar pattern in the human retina to the mice ones (Supplementary Figure 1). Thus, we devised a strategy to profile cells from the retina, as illustrated in Figure 1A. Specifically, the retina was first separated into two regions, the macular/foveal and the peripheral. For the macular/foveal region, as cell proportion was relatively even, we directly profiled nuclei without enrichment. In contrast, for the retinal peripheral region, nuclei are fractioned based on the NeuN staining. As shown in Supplementary Figure 1B, nuclei with high NeuN signal (top 5%, denoted as NeuNT) and moderately high NeuN signal (from 5% to 10%, denoted as NeuNM) are collected for snRNA profiling. The NeuNT group showed the highest enrichment of RGCs and ACs and the NeuNM fraction is enriched of ACs but not RGCs compared to the unenriched peripheral retina sample (Supplementary Figure 1C). Given that snRNA-seq, instead of snATAC-seq, is the major approach for us to perform the cell type classification, we decided to only collect snATAC-seq data from the macula region which already has a relatively balanced cell type distribution (Figure 1A).

**Figure 1.**
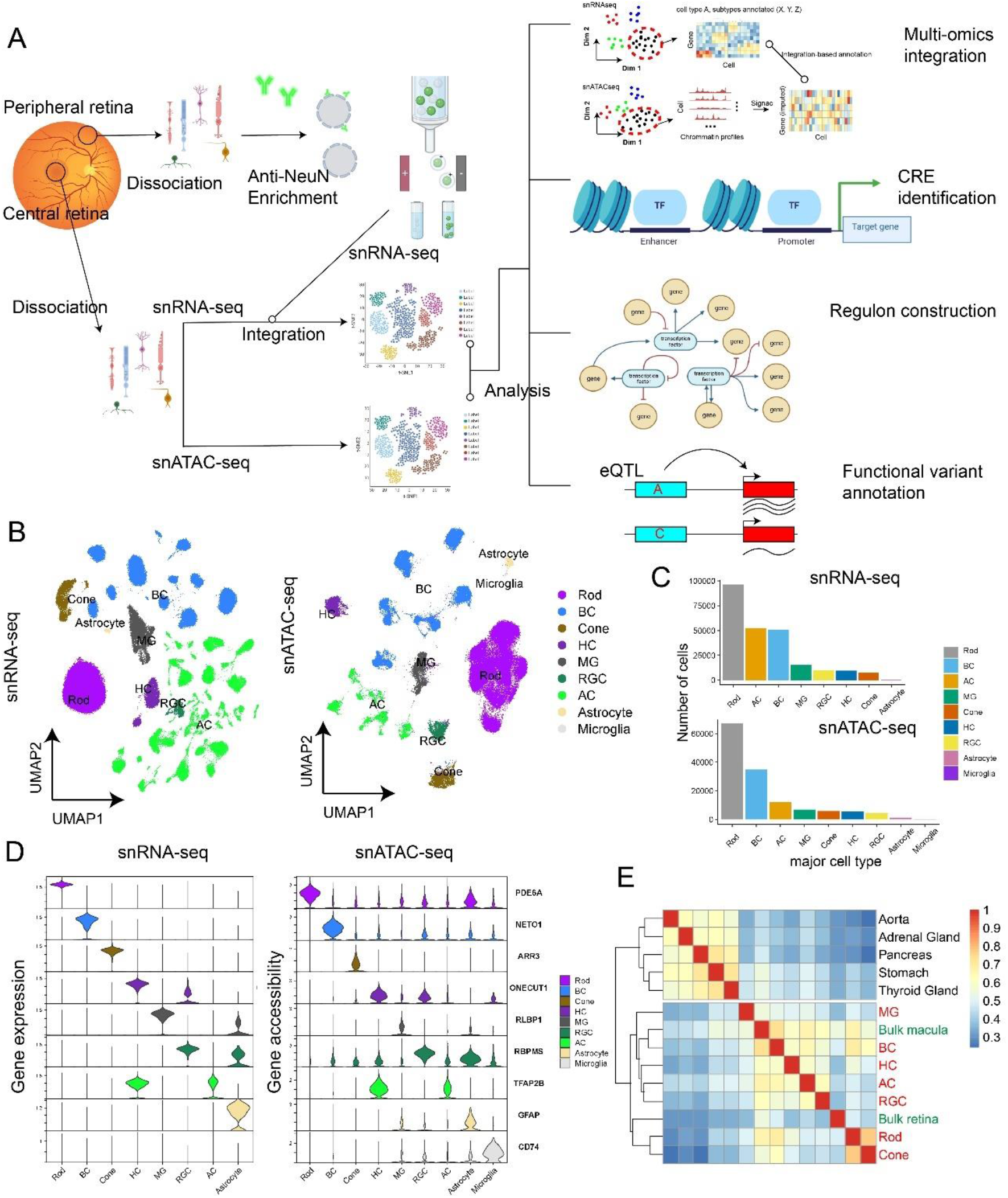
Overview of the single-cell multi-omics atlas of human adult retina. A. The study design of this work. The retina samples were first split into the central and peripheral parts and rare cell enrichment was performed for peripheral retina. The snRNA-seq and snATAC-seq data were first processed separately and then integrated for analysis. B. Two-dimensional embeddings (UMAP) for snRNA-seq (left) and snATAC-seq (right) data. Each data point represents a cell, and the color represents the annotated cell class. C. The number of each cell class from the snRNA-seq (top) and snATAC-seq (bottom). D. The gene expression and gene accessibility of reported retinal cell class marker genes in the snRNA-seq and snATAC-seq data. E. Heatmap of the correlation between the open chromatin profiles of each retinal cell class and those of other human tissues.

We first performed the clustering and cell type annotation on the snRNA-seq and snATAC-seq separately (Figure 1B-C) and all the major cell types showed well isolated, tight clusters. In addition, cluster heterogeneity is visible for some cell classes with multiple types, such as ACs and bipolar cells (BCs). For both datasets, rod cells remain the most abundant major type (Figure 1C), while we were also able to profile over 50k ACs and BCs, and 10k RGCs. Known markers were used to perform the cell type annotation and as is shown in Figure 1D, we visualized the expression and gene body accessibility of the same set of the markers (PDE6A for rod, NETO1 for BC, ARR3 for cone, ONECUT1 for HC, RLBP1 for MG, RBPMS for RGC, TFAP2B for AC, GFAP for astrocyte and CD74 for microglia). In both datasets, all the markers showed quite distinct and expected distribution among cell types. To further evaluate the data quality of the snATAC-seq data, we split the data based on major cell types and compare them with published bulk ATAC-seq data (Figure 1E) of human tissues, including aorta, adrenal gland, pancreas, stomach, thyroid gland, macula, and retina^28^. The datasets formed two groups, with one exclusively for the bulk and single-cell retina data and the other for all other tissues. Moreover, the bulk retina clustered closer to the photoreceptor cells while the bulk macula closer to interneurons, which is expected and likely driven by the different cell type proportion of the bulk and macula samples.

### Transcriptomic and open chromatin profiles of retinal cell types

To obtain the profiles for both the transcriptome and open chromatin modality for retinal cell types, we adapt a strategy that we first annotate types using the RNA-seq data solely, because of the larger number of cells, and then use the annotated RNA-seq data as a reference to annotate the ATAC-seq data (Figure 2A). To determine the optimal parameters for clustering, combination of two parameters, the ‘number of nearest neighbors’ and the ‘Leiden resolution’, that gives the highest Silhouette score is identified (Supplementary Figure 2). As a result, we identified 36 ACs, 14 BCs, 2 HCs, 13 RGCs, 3 photoreceptors (rod and 2 cone types), and 2 non-neuronal cells (MG and astrocyte) in our snRNA-seq data, totaling 70 cell types (Supplementary Figure 3). We then compared this classification result with a previously published human retina transcriptome atlas^9^. BCs and HCs showed high consistency between the two independent datasets with adjusted-rand-index (ARI) close to 1, while ACs showed some differences (we detected 10 more AC subtypes in the current dataset). Through the cross-dataset comparison, we are able to identify several well-studied AC types, such as the starburst AC (AC7), the CAI/CAII ACs (AC34), and VG3 AC (AC28), which all showed one-to-one matching. RGCs have the most differences (Supplementary Figure 4). This is likely due to the fact that both datasets are under powered and have reached sufficient coverage for the rarest RGC types. Nevertheless, two OPN4 positive clusters in our dataset showed one-to-one matching to the two ipRGC clusters of the previous study, and their marker genes, EOMES and LMO2, were also consistent (Supplementary Figure 5). It has been estimated that ipRGCs accounted for around 1% of all the RGCs, which sets the limit of resolution in our presented dataset.

**Figure 2.**
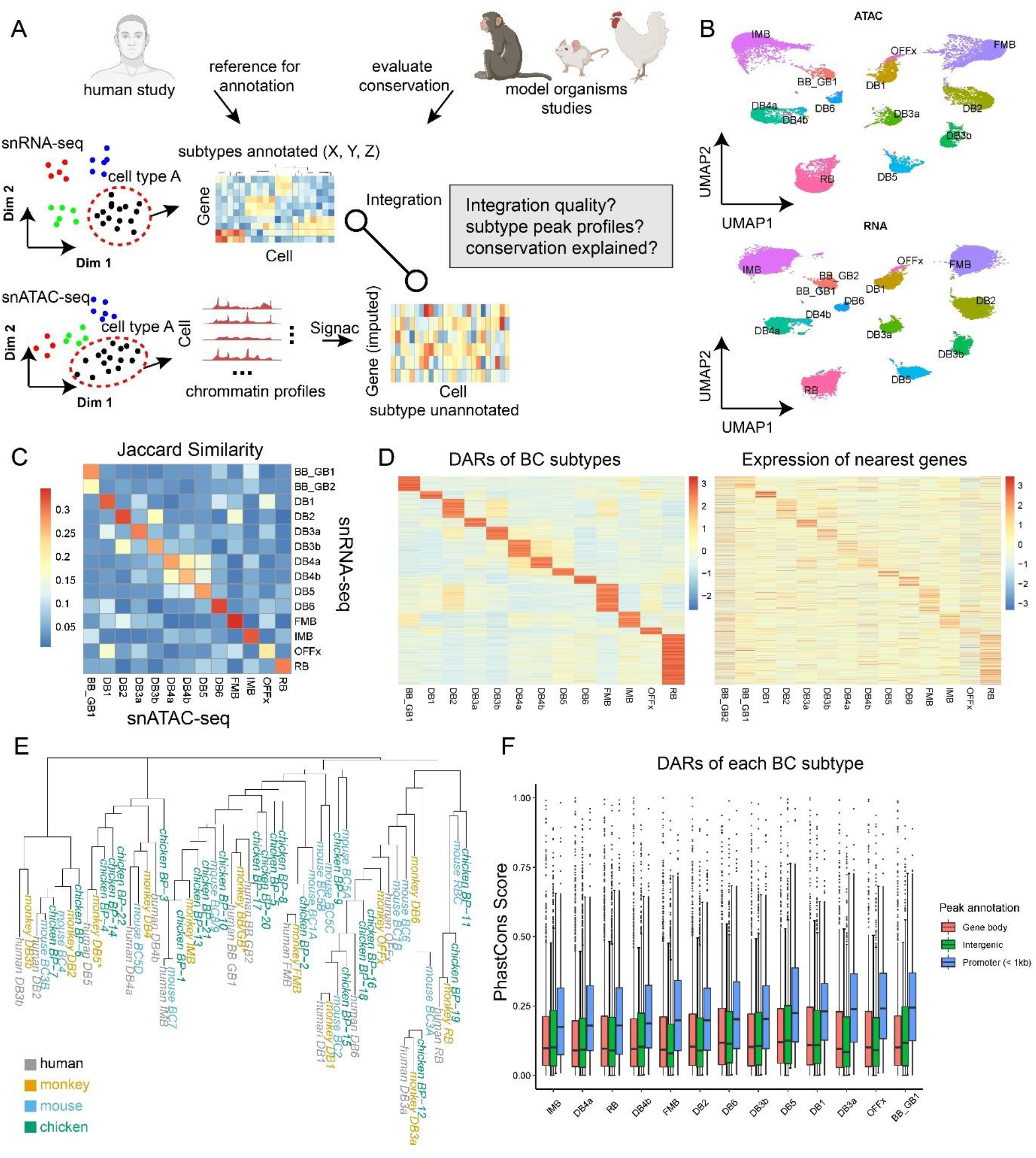
Classification and multi-omics integration of retinal cell types. A. The analysis strategy to perform classification and multi-omics integration of retinal cell types. We perform sub-clustering solely using snRNA-seq data first and leverage published single-cell RNA-seq data of human or model organisms to assist the annotation of the cell types. We then annotate the snATAC-seq data using the annotated snRNA-seq data as the reference. B. Two-dimensional embeddings (UMAP) for bipolar cells of the ATAC (top) and RNA (bottom) modality. C. Heatmap representing the similarity between the differentially expressed genes and the differentially accessible genes from each bipolar cell type. D. Demonstration of the consistency between the differentially accessible regions (DARs, left) and their nearest genes (right) among bipolar cell types. E. Phylogenetic tree representing the overall similarity of bipolar cell types among four species: human (grey), monkey (yellow), mouse (blue), and chicken (green). F. Boxplot showing the PhastCons score of DARs of each bipolar cell type. The DARs were partitioned to ‘gene body’, ‘intergenic’, or ‘promoter’ before the visualization.

With the annotated types from the snRNA-seq data, we performed data integration to co-embed the snRNA-seq and snATAC-seq data followed by the label transfer using the Seurat ‘transfer anchors’ framework^29^. As is shown in Figure 2B, BC types were well separated in the co-embedding space, and the predicted labels of the snATAC-seq cells matches well with snRNA-seq clusters. To further evaluate the reliability of this approach, we first identified differentially expressed genes (DEGs) among cell types using the snRNA-seq data and snATAC-seq data (imputed gene expression) separately, and then computed the pairwise Jaccard similarity between the two DEG lists (Figure 2C). The assumption is that if the label prediction is accurate, signatures detected from one modality of a type should be largely consistent with the ones from the other modality. For most types, the two lists generally showed one-to-one matching (Figure 2C, Supplementary Figure 6). We also computed the signature peaks of BC types (differentially accessible regions; DARs) and showed that their nearest genes displayed a similar pattern in the BC snRNA-seq data. Both results supported the reliability of the snATAC-seq annotation of our data. Thus, we obtained the open chromatin profile of the retina at the individual cell type level.

Cross-species comparison between our dataset and the cell atlas of monkey^30^, mouse^6,8,31,32^, and chicken^7^ is performed. Given that no ATAC-seq based study on human or any model organisms has reached type level, only snRNA-seq data is used. We first co-embedded the data of all the four species into the same latent space, followed by clustering them based on their average distances in the space (Figure 2D). Examination of the result suggests accurate co-embedding has been achieved. For example, the BP-19 from the chicken bipolar cells has been annotated as the chicken rod bipolar cell (RBC). In our phylogenetic tree, this type formed a cluster with the RBCs of the other three species, indicating that the RBC is a highly conserved type, while this BP-19 clustered together with human and monkey flat midget bipolar cells (FMBs) in the previous study^7^. We observed that human and monkey BCs and HCs generally formed one-to-one matching, while ACs show more complex pattern (Figure 2E, Supplementary Figure 7). It is worth noting that even for BCs, some interesting matching patterns are observed. For example, human DB3b and DB5 clustered only with the monkey counterpart, indicating that they could be primate specific types. Similarly, some mouse or chicken bipolar cell clusters are without closely related cluster from human and monkey, indicating they are likely to be species specificity as well. We were then interested that whether the relative conservation of cell types is related to the sequence conservation of their signature genes. We computed the PhastCons scores^33^, an evaluation of the sequence conservation, for each of the DARs of bipolar cell types, and then visualized the scores of the DARs after partitioning them based on the genome distribution: gene body, promoter region, and intergenic regions (Figure 2F). The sequence conservation in promoter regions is constantly higher than those of the other regions. Interestingly, it was not observed that highly conserved types, such as RBCs, showed a higher sequence conservation score in DAR. This result suggested that activity rather than the sequence of the regulatory elements drives the difference among cell types.

### Identification of cis-regulatory elements through integrative analysis of snRNA-seq and snATAC-seq data

The integration of the snRNA-seq and snATAC-seq allows the imputation of gene expression levels for each cell in the snATAC-seq data. Thus, we leveraged this information to compute the correlation between each gene and relatively proximal peaks (Figure 3A). For each gene, open chromatin regions detected by snATAC-seq within 250 kb from the gene transcription start site are tested (TSS). With a cutoff at 0.3 for the Pearson’s correlation coefficient, over 35,000 peak-gene pairs were identified as putative enhancers (Supplementary Table 1, Figure 3B), and they generally showed a closer distance to the gene TSS and a higher sequence conservation (Figure 3C). We also compared the putative enhancer list with the major cell type DARs and found that the list enriches DARs compared to other proximal peaks (within 250 kb) (Figure 3D). Also, as is shown in Figure 3E, both these putative enhancers and their linked genes displayed high levels of cell type specificity.

**Figure 3.**
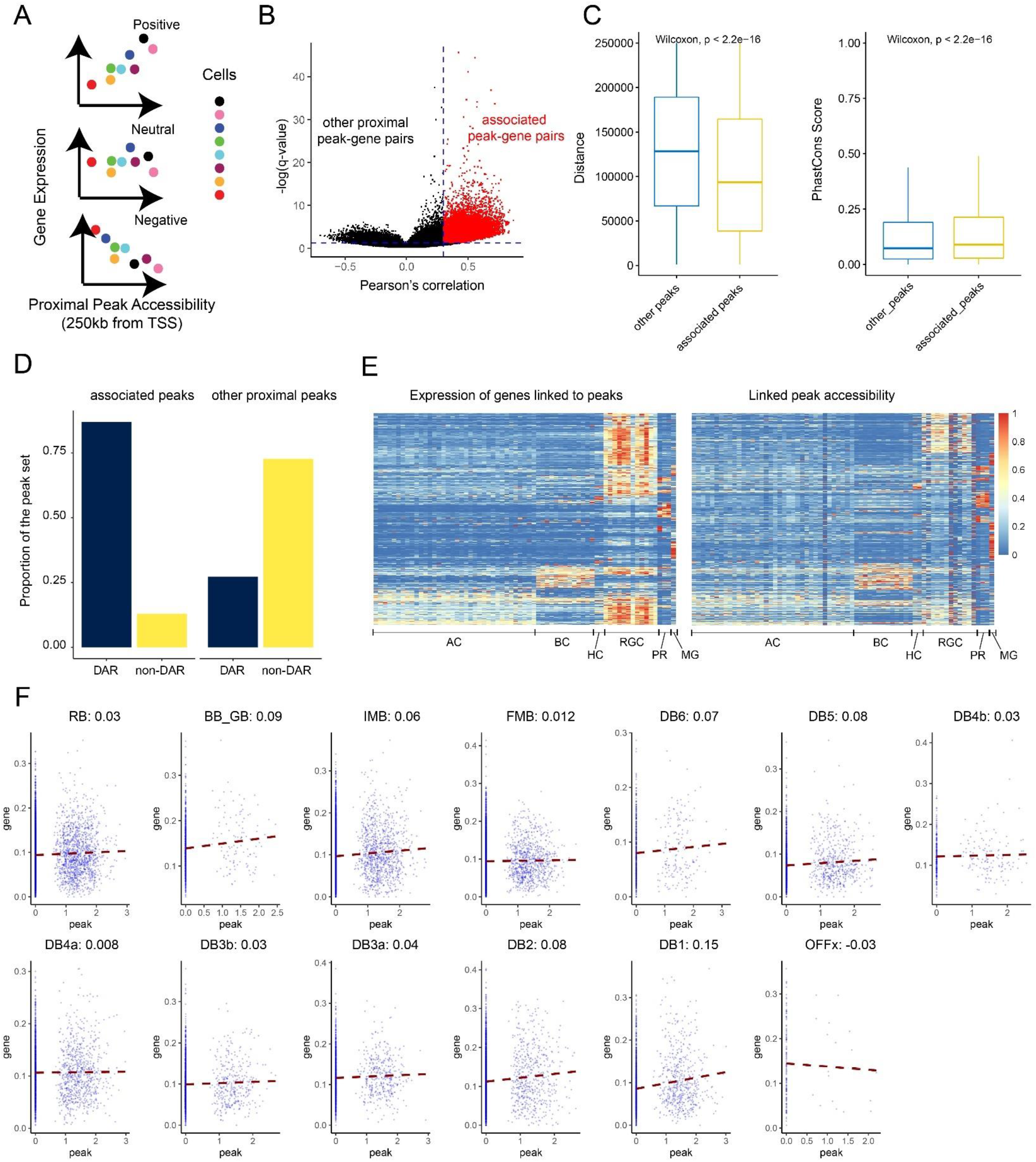
Identification of putative enhancers by linking peak to genes. A. Demonstration of the idea of correlating the gene expression with the proximal peak accessibility. When observing the relationship between a peak and a gene among cells, it is possible that they are positively, negatively, or neutrally correlated. B. Volcano plot showing the overall distribution of peak-gene links in two dimensions: their Pearson’s correlation coefficient and negative log-transformed q-values (p-values with FDR correction). Red data points highlight the links we considered as putative enhancers. C. Boxplot showing the comparison between the putative enhancers and other proximal peaks on their distance to linked gene and sequence conservation. Wilcoxon tests were performed to evaluate the difference between the groups. D. Bar plot showing the comparison between the putative enhancers and other proximal peaks on their overlapping with major cell class DARs. E. Heatmap showing the relative expression of genes linked to peaks and their linked genes across different cell types. The gene expression level and peak accessibility level were both normalized between 0 and 1. F. The peak gene correlation between chr21:45358487-45360141 and CSTB in all bipolar cell types. For each panel, each data point represents a cell and the red dashed-line represents the linear fit of the correlation between the peak and the gene. The Pearson’s correlation coefficients were labeled on each panel.

Correlating peak accessibility to gene expression across cells has been incorporated in different analysis frameworks as a method to identify regulatory regions^34,35^. However, it has not been well discussed about what groups of cells should be used as the input. A simple question could be whether to use all the cells as the input, or only use cells from the same cell type as the input. To obtain a core set of putative enhancers, we used all the cells as the input and we reason that this approach will allow for detecting stronger signals, since a much larger number of cells are used. However, given the complexity of the retinal types, we hypothesized that while some peaks have consistent activities, other peaks might display different or even opposite activities among cell types of the same class. Thus, we performed the peak-gene correlation analysis for each BC types separately. For each peak-gene pair, the mean and standard deviation of the Pearson’s correlation coefficient across the cell types is calculated (Supplementary Figure 8A). Both constant and highly heterogenous peak-gene pairs have been identified. As an example of a constant peak-gene link (chr21:45358487-45360141; CSTB; Figure 3F), all the BC types except OFFx showed a positive correlation. As an example of heterogenous link (chr12: 56614952-56617057; ESYT1; Supplementary Figure 8B), six BC types showed negative correlation while the rest showed positive correlation. Admittedly, the correlation coefficient looked less significant, which we reason is caused by the high sparsity of the single-cell data and remained a huge challenge for the field. With these results, we showed cases that the activity of CREs could differ among cell types of the same cell class, therefore underscores the importance of obtaining high resolution chromatin profiling.

### Integrative analysis of transcription factor functions and cell type specificity in the retina

We then inferred transcription factor activities for retina cells leveraging both the transcriptome and open chromatin information (Supplementary Figure 9). We first created ‘meta cells’ for each type based on their similarity in latent space (20 cells into 1 meta cell) and used the meta cells as the input. Our approach was based on the SCENIC pipeline^36^, with a modification that we trimmed the initial TF-target regulon based on the appearance of the TF motif in proximal peaks of the potential target gene (Supplementary Figure 9). We then scored the TF activity for each meta cell using the AUCell method (Figure 4A). It could be observed that most of the TFs (200) showed certain levels of cell type specificity (Figure 4A). The number of target genes differed drastically across the TFs, with a median of 196 targets per TF. For each target gene, the median of the number of TFs was 6 (Figure 4B). The full TF-target list could be found in Supplementary Table 2.

**Figure 4.**
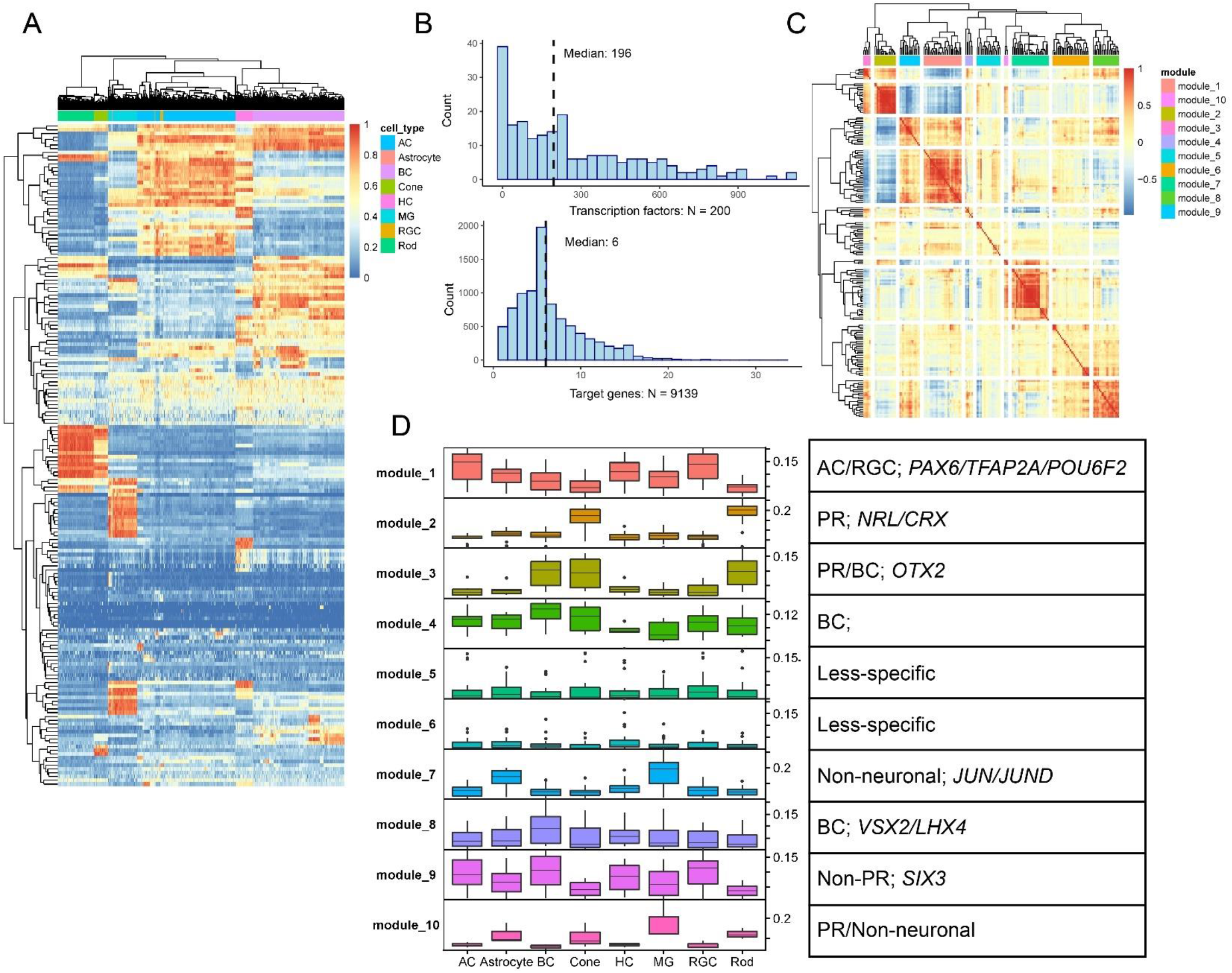
Inference of TF activity and annotation of TF modules. A. Heatmap representing the TF activities (row) for each meta cell (column). The TF activities were first calculated using the AUCell framework and then normalized between 0 and 1. B. Basic metrics of the TF-target regulons. The upper and bottom panels showed the distribution of number of targets per TF and numbers of TF regulators per target. Median values were labeled. C. Heatmap depicting the TF-TF correlations. Hierarchical clustering was first made, and modules were detected by cutting the tree to K groups (K = 10 here). D. The activity of TFs in each module in each cell classes. The TF modules were then annotated based on the observed specificity of the module. Previously studied TFs were also listed.

We then computed the correlation between TFs using their activity as the input (Figure 4C) and grouped them into 10 modules. Each TF module is annotated based on their activity in classes (Figure 4D). We found that rather than being specific at individual cell class level, most of the TF modules accounted for combinations of cell classes, such as the ones for photoreceptor-bipolar (module 3), AC-RGC (module 1), non-photoreceptor (module 9), etc. There were also two modules (5/6) being less specific. The full list of the partition of TFs modules and the TF activity at major cell class level could be found in Supplementary Table 3. Finally, we investigated the TF-TF regulatory relationships by building a directed network for the TFs. The out degree is a measurement of the number of direct targets for a TF in this network, and we reason that the TFs with high out degree could be potential hub regulators (Supplementary Figure 10A). We selected the TFs with out-degree larger than 15 and calculated the overlapping between their targets and the TFs in each of the 10 modules (Supplementary Table 4 and Supplementary Figure 10B). Some well-studied TFs were identified. For example, PAX6 appeared as a potential hub regulator for module 1, which was the AC/RGC module. Also, CRX, OTX2, and NRL appeared as potential hub regulators for module 2, which was the photoreceptor module. These examples highlighted the reliability of the analysis, while other less studied TFs still required further study.

## Discussion

### Comprehensive single-cell multi-omics atlas of the human retina

In this study, we have generated a comprehensive single-cell multi-omics atlas of the human retina that includes snRNA-seq for over 250K nuclei and snATAC-seq for over 137K nuclei. To capture the rare cell types in the retina, we have sorted nuclei based on NeuN gradient and collected fractions that are enriched for ACs and RGCs. This combined strategy resulted in an atlas that is significantly improved over previous published dataset in several aspects. First, the cell portion is better normalized for cell types with high heterogeneity, including AC, BC, and RGC, comprising 21.5%, 20.8%, 4.1% of the cell population for this dataset respectively. Consequently, this allows detection of rare cell types, such as the two ipRGC types, which are estimated to comprise only 0.01% of all retinal cells. Second, the number of cell types identified in this study is 69, including 67 neuronal types, which is the highest reported among all published studies. This likely to represent a near complete catalogue of all cell types in the retina. Third, this study is further strengthened with the first large-scale snATAC-seq dataset in the retina with 137K nuclei profiled. Admittedly, the detection of RGCs was still underpowered in our dataset. This is possibly because we only applied enrichment using the peripheral retinal samples, and ACs were the more dominant NeuN positive cell class, not RGCs. A full investigation of RGCs would need further efforts. This data set represents the most comprehensive and information rich resource for understanding retinal function and pathology at the cellular level.

### Cross-species comparison among cell types

As the key light sensing organ, the evolution of the eye is fascinating and the morphology and structure of the eye across metazoan is highly diverse. Vertebrate animals share the camera like eye structure which contains a multi-layer neuronal structure called retina whose function is to capture the light and covert it to electric signal that is transmitted to brain. The major cell types in the retina, which includes photoreceptor, bipolar, horizontal, amacrine, and retinal ganglion cell, are conserved across vertebrate animals. With the comprehensive retina cell atlas in hand, it is possible to conduct cross-species comparisons among human, monkey, chicken, and mouse at cell type level. Interestingly, we found that the degree of conservation differs among different cell type with increased divergence observed as we move from the out nuclear to the inner nuclear layer of the retina. Specifically, conservation decreased from outer to inner retina, with cell types at the outer nuclear layer of the retina being the most conserved and cell types in the retinal ganglion layer being the most divergent. This finding is consistent with a previous study^5^ and suggests that although there is a common mechanism of initial retinal photon capture, downstream processing of the visual signal diverges between species. Interestingly, the number of cell types observed in ACs and RGCs is negatively correlated with the position of the organism in the evolution tree. Much large number of AC and RGC types have been identified in mice and chicken retina compared with human and monkey. A potential model to explain this phenomenon is that higher animals might rely more on the visual cortex for processing visual signals.

Our approaches to perform the comparison is also worth mentioning. There has not been a standard, widely applied workflow designed aiming at computing the distance between cross-species cell types from scRNA-seq data. Regardless of which metric was selected, direct measurement of distance among cross-species cell types tended to group the cell types from the same species together. This was due to significant ‘batch effects’, which were stronger than normal batch effects driven by different sequencing technologies or by different groups that generated the data. Thus, we chose to first minimize the batch effects across the datasets through data integration using LIGER^37^, which performed the best out of several methods, many of which failed in this type of task, according to Luecken et al^38^. The distance among cell types was then computed in the reduced dimensions after integration. This approach allows us to capture the conserved signal relatively accurately across species, such as the chicken BP-19 case as mentioned above.

With multi-omics data at the cell type level, we were able to test hypothesis regarding the relationship between gene regulation landscapes and the conservation of subtypes. The first natural hypothesis we generated was that the regulatory elements for signature genes of conserved subtypes might also be more conserved. We found that the conservation of promoters of these genes could not explain the conservation of subtypes, indicating the involvement of more complex regulatory processes.

### Integrative analysis for snRNA-seq and snATAC-seq to decipher key elements in gene regulation

Cis-regulatory elements (CREs) are important genomic regions that regulate the transcription of neighboring genes usually through regulating the recruitment of transcription initiation machinery. With the two modalities of gene expression and peak accessibility, we associated them to infer the potential regulatory functions of genomic regions. Over 35k peak-gene pairs were identified in our study and most of the peaks were also identified as DARs for at least one cell class. Besides, we investigated the regulatory function of genomic regions in different types within a major type. We showed that some genomic regions could constantly act as regulators for proximal genes, while others displayed higher heterogeneity. This finding highlights the importance of investigating CREs in the context of cell types. Also, this could only be investigated with a comprehensive dataset as we presented here, for cell type level resolution is required for both modalities.

In addition to identification of CREs, mapping chromatin accessibility can also help to study gene regulatory networks (GRNs). Many efforts have been made, using scRNA-seq alone, to decipher the GRNs for transcription factors or simply build gene modules. One of the popular frameworks, SCENIC^36^, used the motif information at the proximal regions of genes (e.g., 10kb regions around TSS) for trimming the initially built GRNs. With the snATAC-seq data, the search space for motif existence become broader and more accurate. In this study, we leveraged the information of the TF-motif presence in open chromatin regions around gene TSS to perform the pruning of co-expression-based TF-target networks. With this approach, we were able to quantitatively assess the activity of TFs and learn their specificity across cell types. Furthermore, we could group the TFs into modules based on their activity in cells and annotate these modules. Multiple well-studied cell type specific TFs such as OTX2, PAX6, and NRL were included in the expected modules and were likely the key regulators for these modules.

In summary, we report the first comprehensive single-cell multi-omics atlas for the human adult retina as part of the HCA project. The utilization of well characterized retinal tissue in this study increases the translational value of the resulting data set. This atlas enables in-depth integrative analysis at individual cell type resolution, making it a highly valuable and robust resource for the research community.

## Methods

### Human retina sample collection

The samples used for this study were collected from 25 individuals from the Utah Lions Eye Bank. Detailed information for these donors was included in Tables 1 and 2. All donor eye samples were collected within 6 hours post-mortem. Donor eyes were subsequently phenotyped to ensure that they were absent of any disease pathology. Details for the post mortem phenotyping and dissection were according to a previously described standardized protocol^39^. In brief, only samples from donors with no record of diabetes, retinal degeneration, macular degeneration, or any other retinal diseases were used in this study. Post-mortem phenotyping with spectral domain –optical coherence tomography (SD-OCT) and fundus imaging, to confirm that there were no drusen, atrophy, or other disease pathology, was done utilizing our standardized approach^40^. Although *both* donor eyes were phenotyped for each subject, to ensure that disease pathology was absent for a given subject, only one donor eye was used. For this study, the fovea and macula samples were collected separately, using a 4 mm and 6 mm disposable biopsy punch, respectively, and were flash-frozen in liquid nitrogen. All samples were then stored at −80 °C before nuclei isolation.

All tissues were de-identified under HIPAA Privacy Rules. Institutional approval for the consent of patients for their tissue donation was obtained from the University of Utah and conformed to the tenets of the Declaration of Helsinki.

### Nuclei isolation and sorting

Nuclei were isolated by pre-chilled fresh-made RNase-free lysis buffer (10 mM Tris-HCl, 10 mM NaCl, 3 mM MgCl2, 0.02% NP40). The frozen tissue was resuspended and triturated to break the tissue structure in lysis buffer and homogenized with a Wheaton™ Dounce Tissue Grinder. Isolated nuclei were stained with mouse anti-NeuN monoclonal antibody (1:5000, Alexa Flour 488 Conjugate MAB377X, Millipore, Billerica, Massachusetts, United States) in pre-chilled fresh-made wash buffer (1% BSA in PBS, 0.2U/μl RNAse inhibitor) for 30min at 4°C. After being centrifuged at 500g, the pellet was resuspended in wash buffer and filtered with 40μm Flowmi Cell Strainer. DAPI (4′,6-diamidino-2-phenylindole, 10 μg/ml) was added before loading the nuclei for fluorescent cytometry sorting.

Stained nuclei were sorted with a FACS Aria II flow sorter (Becton Dickinson, San Jose, CA), (70μm nozzle). Sorting gates were based on flow analysis of events and strengths of DAPI signal, as well as FITC signal. Samples were sorted at a rate of 50 events per second, based on side scatter (threshold value >200). Fluorescence detection used a 450-nm/40-nm-band pass barrier filter for DAPI, and a 530-nm/30-nm-band pass filter for FITC. For the NeuNT group, the nuclei with the strongest 5% FITC signal were collected into Eppendorf tubes with 3μl pre-chilled wash buffer.

### Single-nuclei sequencing

All single-nuclei RNA or ATAC sequencing in this study was performed at the Single Cell Genomics Core at Baylor College of Medicine. Single-nuclei cDNA library preparation and sequencing were performed following manufacturer’s protocols (https://www.10xgenomics.com). Single-nuclei suspension was loaded on a Chromium controller to obtain single cell GEMS (Gel Beads-In-Emulsions) for the reaction. The snRNA-seq library was prepared with Chromium Next GEM Single Cell 3’ Kit v3.1 (10x Genomics). The snATAC-seq library was prepared with Chromium Next GEM Single Cell ATAC Library & Gel Bead Kit v1.1 (10x Genomics). The library was then sequenced on Illumina Novaseq 6000 (https://www.illumina.com).

### Sequencing read alignment

Reads from snRNA-seq were demultiplexed and then aligned to the ‘Ref_cellranger.hg19.premrna’ human reference (from 10x Genomics) using Cell Ranger (version 3.0.2, 10x Genomics). Reads from snATAC-seq were demultiplexed and then aligned to the refdata-cellranger-atac-hg19-1.2.0 human reference (from 10x Genomics) using Cell Ranger (version 1.2.0, 10x Genomics).

### snRNA-seq quality control, clustering, and differential expression

The gene expression matrices generated by Cell Ranger were filtered for quality control purposes. We first used the soupX package^41^ to correct the feature counts to alleviate the effect of ambient RNA. For each matrix, genes detected in less than 5 cells were removed; cells with total UMI counts less than 800 or more than 8000 were removed, and top 5% of cells with highest mitochondrial gene expression proportion were removed. This processing was performed using the Seurat R package^42^. Unlike single-cell RNA-seq, snRNA-seq data had a much lower mitochondrial gene expression proportion, and thus, we removed high-mitochondrial-content cells based on the rank of cells but not a fixed threshold of mitochondrial gene expression proportion. We then performed doublet removal on the remaining cells using DoubletFinder^43^. The key parameters of DoubletFinder were set such that pN equals 0.25, the estimated doublet ratio was 0.1, and pK was estimated automatically with ‘find.pK’ function.

The expression matrices after the initial QC were combined to one single scanpy^44^ anndata object. To account for batch effects originating from sample collection processes, the scVI framework^45^ was applied with standard parameters. The low dimensional embeddings generated were then used to perform leiden clustering^46^, and major cell classes were annotated based on known marker genes. Major cell classes were then split for sub-clustering, which were also performed with scVI with the same process as described above. To avoid under-or over-clustering in type discovery, we adapted a parameter searching approach to find the best combination of neighbors and resolution for clustering, as described in Supplementary Figure 2. After sub-clustering, we checked the expression of marker genes for all major cell classes and used that to remove doublets for a second round. For example, if an AC cluster had high rod marker gene expression, it was removed.

For the differential expression analysis, we used the default Wilcoxon method implemented in the Seurat package. The ‘logfc.threshold’ was set to 0.25 and ‘min.pct’ set to 0.2. Only genes with an adjusted p-value less than 0.05 were retained as differentially expressed genes (DEGs). The DEGs for major cell classes were computed using other classes as background, while the DEGs for types were computed using other types from the same class as background.

### Obtaining additional RGC data from other local datasets

To increase the number of RGCs for building a more comprehensive atlas, besides the cells from the six donors primarily used, we also used RGCs from other locally sequenced datasets. These RGCs were identified using a supervised single-cell annotation framework, scPred^47^. Annotated snRNA-seq data, including rod, cone, MG, AC, BC, HC, and RGC, were downsampled to 10,000 cells per cell type, as input for training the scPred model. 70% of cells were used for training and the rest were used for evaluating the performance of the model. The model was then applied to query datasets for automatically annotating cells to major classes, with the prediction threshold set to 0.8.

### Matching cell types between the current study and previous ones using random forest classifiers

Random forest based classifiers had been shown to perform well in previous primate retina studies^5^. To match cell types between the current study and a previous human retina study, the top 2000 highly variable genes were selected from the human expression matrix and were used to build a classifier on the monkey expression matrix. 500 cells for each cell type were used to train the classifier and the rest were used to evaluate the classifier. Cell types that had small numbers of cells were split by 70% and 30% as training and test set. The random forest classifier was built using the ‘randomForest’ package in R, using ‘ntree’ equal to 1001. For each cell, if the highest prediction voting rate was less than a threshold (20%), it was labeled as ‘unassigned’. Sankey plot was used to visualize the matching of the data, performed in python with the plotly.graph_objects package.

### Cross-species integration of snRNA-seq data

The expression matrices and annotations for single-cell RNA-seq data for mouse, chicken, and monkey retina were obtained from previous publications^5,7,8,31^. All the gene names were converted to human gene nomenclatures, and the matrices from the three species were then merged to a single matrix, for each cell type. At this step, we only retained the genes with 1:1 matching orthologs across species. The data were then integrated using the LIGER framework^37^, implemented as a Seurat wrapper. A standard workflow was applied, using the top 5000 highly variable genes, with k value set to 20 and lambda set to 5. The data was projected to an integration space with 20 dimensions. The coordinates of cells from the same type were averaged and then used to calculate the distance matrix (across each species) using the ‘Dist’ function of the ‘amap’ R package^48^. Euclidean distance was used. The distance matrix was then used to build the phylogeny tree using the ‘nj’ function of the ‘ape’ R package^49^.

### snATAC-seq quality control, clustering, and differential accessible analysis

The snATAC-seq data was mainly analyzed using the Signac^34^ framework. The data output from Cell Ranger ATAC were used as the input. Data from each donor were first combined based on a consensus set of genomic regions and then filtered with QC metrics after merging to a single object. Cells with less than 3000 fragments or more than 20000 fragments were removed for downstream analysis. We also used another two QC criteria implemented in Signac, nucleosome signal and TSS enrichment for filtering nuclei (nucleosome signal less than 4 and TSS enrichment larger than 2 for the nuclei to be retained). Thus, we resulted in an initial peak-by-nuclei matrix.

Dimensional reduction was performed using the ‘RunTFIDF’ and ‘RunSVD’ functions with default parameters and then clustered using ‘FindClusters’ function, also with default parameters. The clusters were annotated to major cell types by checking the genome accessibility of known marker genes. Also, with the annotation information, peaks were identified with macs2^50^ called by each cell class. With the macs2 output, we updated the peak-by-nuclei matrix, and all the downstream analysis were based on this updated version.

Differential accessible analysis for finding differential peaks were performed with FindAllMarkers of the Seurat package, setting the minimal detection percentage (min.pct) equal to 0.1. For calculating the conservation of each peak, the peak coordinates were first converted to a GenomicRanges object, and we used the R package GenomicScores^51^ to score the conservation of each peak using the ‘phastCons 100 species’ system.

### Comparing snATAC-seq data with bulk ATAC-seq data

Public ATAC-seq data in bigwig format were downloaded from ENCODE portal^52^ (ENCFF158OVK for pancreas, ENCFF507OEP for aorta, ENCFF111XAE for adrenal gland, ENCFF961DZL for thyroid gland, ENCFF492WGC for stomach), and GEO (GSE137311, for retina and macula). Conversion from hg38 aligned coordinates to hg19 versions was performed using CrossMap. snATAC-seq data (bam files) were split based on cell classes using Pysam (https://github.com/pysam-developers/pysam) based on cell barcodes. The bam files for each cell type were deduplicated using the ‘MarkDuplicates’ function in picard tools (http://broadinstitute.github.io/picard/). They were then converted to bigwig format, normalized by total reads, and scaled to 100 million reads using the ‘genomecov’ function in bedtools^53^. The genome coverage for all the public bulk ATAC-seq data or grouped snATAC-seq data were transformed to a data frame using the ‘multiBigwigSummary’ function of the deeptools software^54^, using a window size of 10 kb. Spearman correlation among samples was computed using ‘plotCorrelation’ function of deeptools.

### Integration of snATAC-seq and snRNA-seq data

For snATAC-seq data, the gene expression was modeled using the ‘GeneActivity’ implemented in Signac framework. For cell classes with known cell type heterogeneity (AC, BC, HC, Cone, and RGC), the integration was performed within that class, to annotate snATAC-seq data to types. Highly variable genes in both snRNA-seq data and GeneActivity matrices of snATAC-seq data were selected and the common ones were used as inputs for data integration. The GeneActivity matrices of snATAC-seq were first used as query data for further imputing the gene expression using the ‘transfer anchors’ method in the Seurat R package, with snRNA-seq as reference. The imputed gene expression matrices are used for the calculation of the peak-gene correlation at single-cell levels. We also directly predict labels for the ATAC-seq data for the purpose of cell type annotation.

### Identification of CREs

The gene expression for each nucleus in the snATAC-seq data was inferred using the Seurat ‘transfer anchors’ method with the snRNA-seq data as the reference. Thus, for each nucleus, we have both the gene expression and peak accessibility information. These two pieces of information were used as inputs to infer the association between genes and peaks. To avoid the effect of the sparsity of the single-cell data, we created meta cells by averaging cells being neighbors in the low dimensional embedding. Each meta cell is an average of 10 different cells and each cell is only merged into a single meta cell (https://github.com/qingnanl/SRAVG). We used the ‘LinkPeaks’ function in the Signac package to calculate the correlation between each gene and any peak within 250 kb to the gene TSS. We first performed the analysis with all the nuclei as input to calculate an overall peak-gene association (Figure 3B-E), and then repeated the analysis with only each type as the input, one at a time (Figure 3F).

### Transcription factor regulon determination and scoring

The initial construction TF regulons required the gene co-expression analysis using the snRNA-seq data. Given the large size of our data, in consideration of computational efficiency, we created meta cells by grouping cells within each cell type and used the average gene expression for this meta cell. The size of each group was 20, so that the size of this meta cell matrix could be reduced to about 5% of the original one. This approach was previously reported by Suo et al^55^.

The meta cell matrix were then used as the input of GINIE3^56^ algorithm to construct an initial TF regulon network. The output of GINIE3 was a weight matrix regarding how likely each TF is a regulator of each target gene. We set the weight cutoff at 0.005 for deciding the TF-target relationship and thus we resulted in an initial TF-target network. We then used the peaks proximal to target TSS, which could be obtained from snATAC-seq data, to prune the network. We searched motifs in peaks which were 50kb upstream to the TSS and required at least one appearance for the TF-target pair to be retained. With the pruned regulons, we used the AUCell R package by the default parameters to calculate the TF activity score for each meta cell.

### Immunofluorescence (IF) Staining

Healthy human donor retinal tissue was dissected and fixed in 4% PFA for 48 h, cryo-protected with 30% sucrose overnight in 4°C, and then embedded in OCT to be flash-frozen. Cryosections from a peripheral region of the human retina with 10 μm thickness were used for the IF.

For the IF, sections were fixed with 4% PFA for 15 min at room temperature, and after being washed with PBS for 3 times, 5min per time, they were blocked for 3 h with blocking buffer (10% normal goat serum in PBS + 0.1% Triton X-100) at room temperature. Sections were then incubated with primary antibodies (anti-NeuN: 1:50, MAB377X, Millipore, Billerica, Massachusetts, United States; anti-RBPMS:1:1000, Novus Biologicals NBP2-20112) diluted in blocking buffer. The sections were washed and stained with species-specific fluorophore-conjugated secondary antibody in blocking buffer (1:500) for 2hrs at room temperature, then washed and stained with DAPI for 10 min. After washing with PBS 3 times, the sections were mounted with AquaMount Slide Mounting Media (Thermo Scientific 13800). IF stained slides were visualized with Zeiss Axio Imager M2m.

## Acknowledgement

This work is supported by Chan-Zuckerburg Foundation Grant CZF2019-002425. Single-nucleus RNA sequencing was performed at the Single Cell Genomics Core at BCM partially supported by NIH shared instrument grants (S10OD018033, S10OD023469, S10OD025240), P30CA125123, P30EY002520, and CPRIT Comprehensive Cancer Epigenomics Core Facility RP200504. The cell sorting experiments are also supported by the Cytometry and Cell Sorting Core at Baylor College of Medicine with funding from the CPRIT Core Facility Support Award (CPRIT-RP180672), the NIH (CA125123 and RR024574) and the assistance of Joel M. Sederstrom. Ascertainment, collection, processing, and phenotyping of donor eye tissue was partially supported by the Macular Degeneration Foundation, Inc. and the Carl Marshall & Mildred Almen Reeves Foundation.

We want to thank Kaitlyn Xiong for offering critical advice.

## Data and code availability

All sequencing data generated in this study is available at: https://data.humancellatlas.org/explore/projects/9c20a245-f2c0-43ae-82c9-2232ec6b594f

HCA portal for this data could be used for interactive visualization: https://cellxgene.cziscience.com/collections/af893e86-8e9f-41f1-a474-ef05359b1fb7

Original code is available in: https://github.com/qingnanl/human-retina-multiomics

More information could be provided by contacting the correspondence authors.

---

**Supplementary Figure 1.**
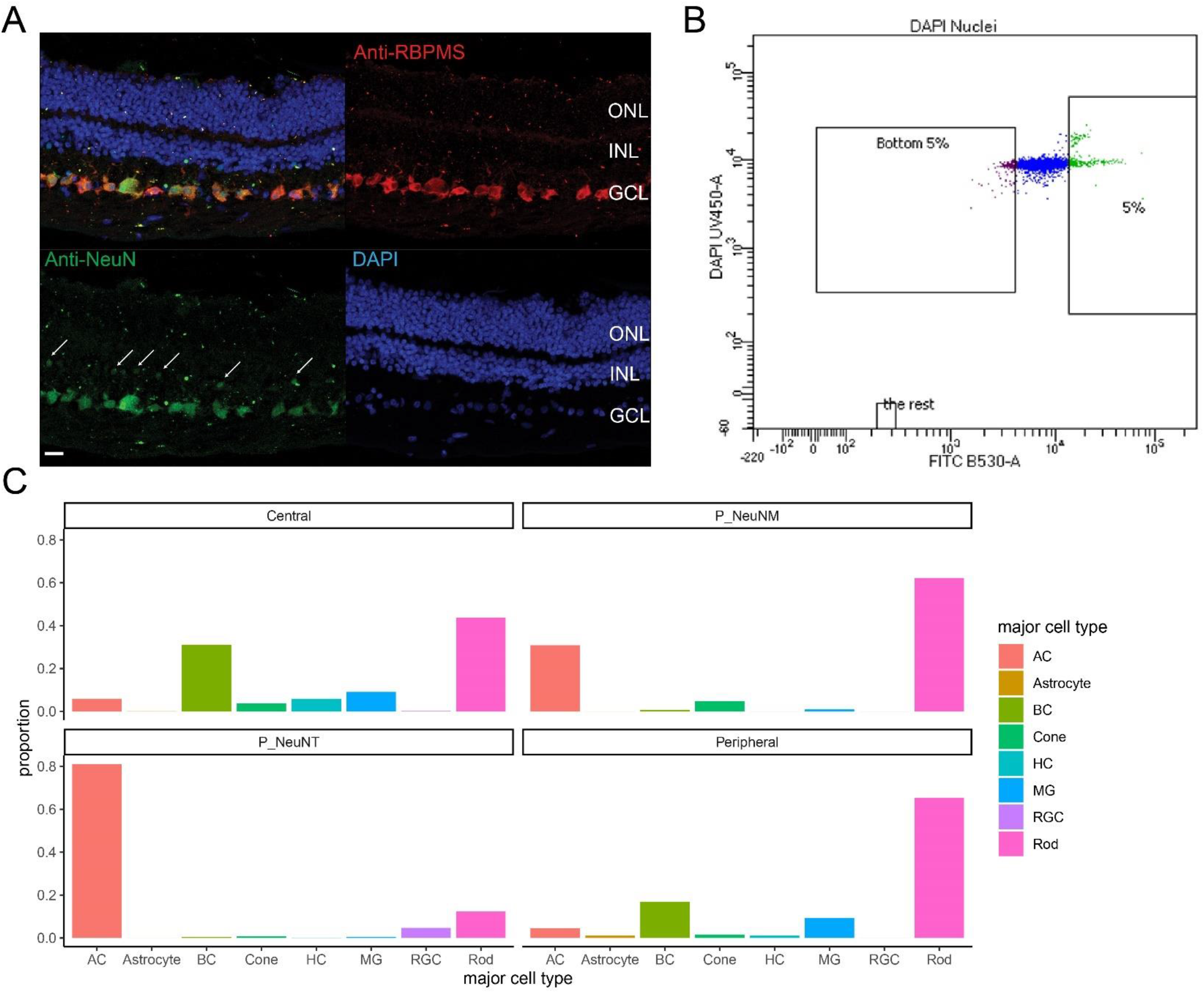
NeuN enrichment of rare retinal cell classes. A. NeuN is highly expressed in the RGCs and amacrine cells in the human retina. Immunofluorescent staining with anti-NeuN (green) and anti-RBPMS (red), a pan RGC marker, was performed on human adult retina sections. The top left showed the merged-channel image with NeuN, RBPMS, and DAPI signals. White arrows highlight NeuN-positive cells in the INL, likely being amacrine cells based on their nuclei location. The scale bar (bottom left) indicates 20 microns. B. Representative FACS sorting plot for separating DAPI positive nuclei with the highest 5% anti-NeuN signal (FITC channel). C. The proportion of major cell classes in four groups of the snRNA-seq data: central retina, peripheral retina with medium-high NeuN signal (P_NeuM), peripheral retina with top NeuN signal (P_NeuNT), peripheral retina with no enrichment.

**Supplementary Figure 2.**
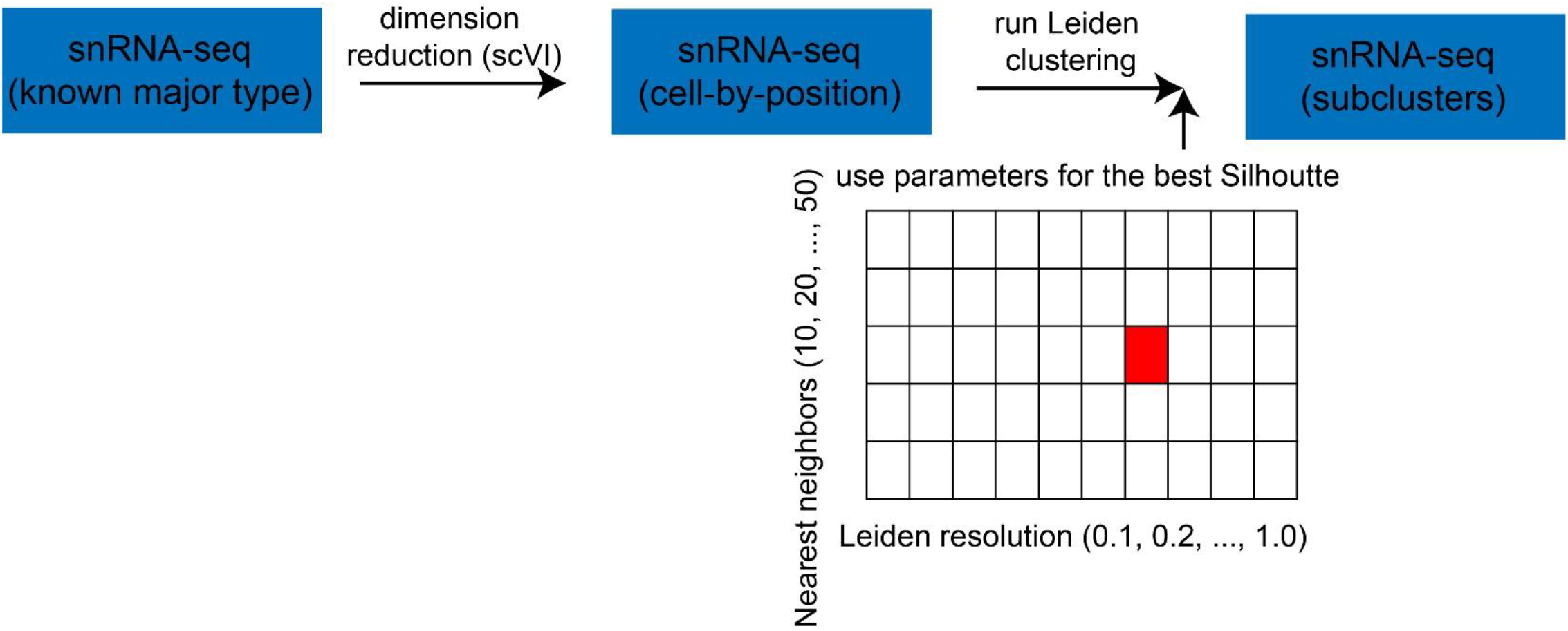
The strategy of detecting cell types. For each complex class (BC, AC, HC, RGC, cone), dimension reduction was performed with the scVI framework. In the latent space, Leiden clustering was performed 50 times with a combination of number of nearest neighbors (10, 20, …, 50) and Leiden resolution (0.1, 0.2, …, 1.0) to find the combination giving the best Silhouette score. The clustering result from that very combination was then used as the optimal clustering used for downstream analysis.

**Supplementary Figure 3.**
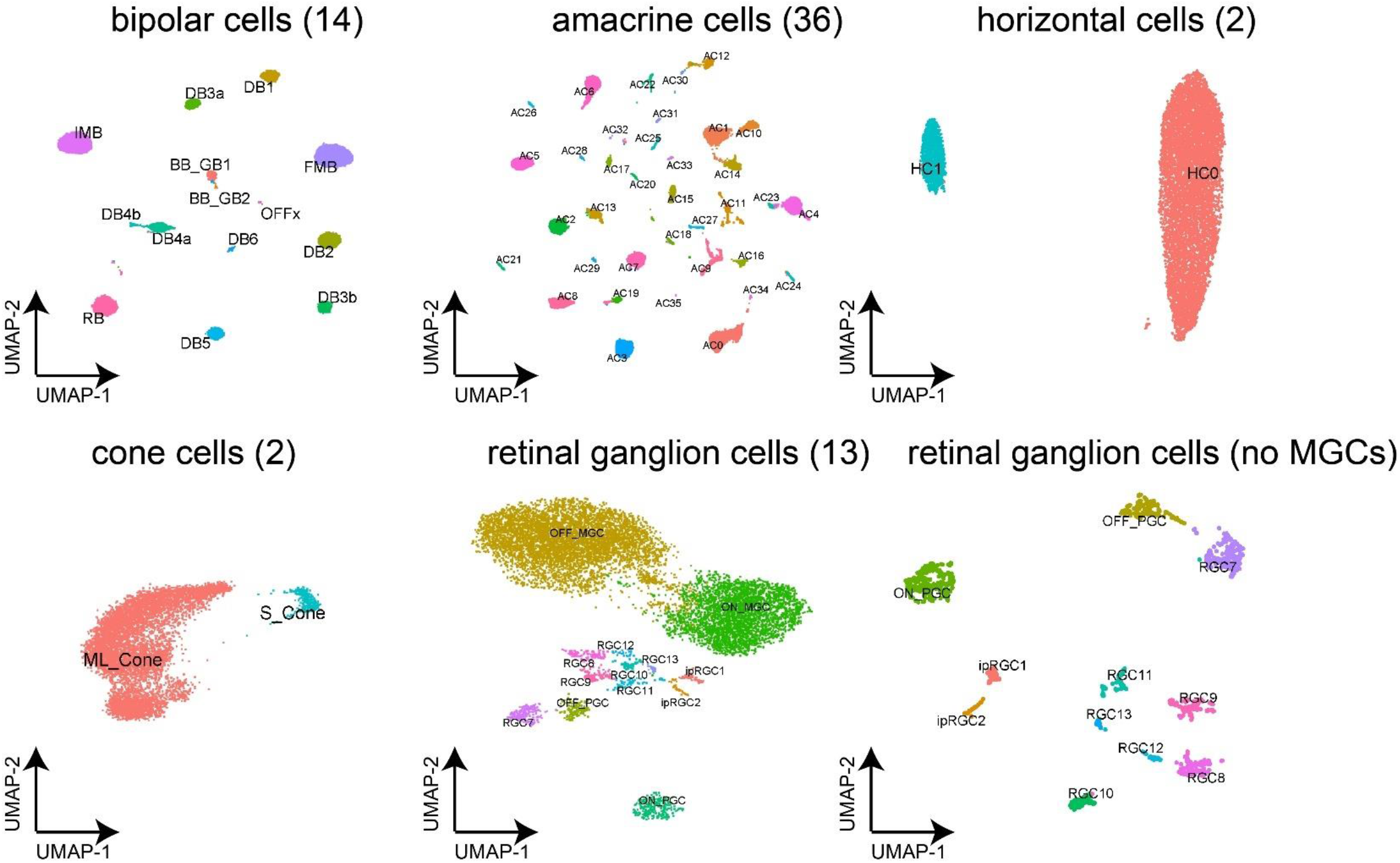
Cell types detected from snRNA-seq data. Two-dimensional embeddings of cell types from BCs, ACs, HCs, cones, and RGCs were visualized. For RGCs, given that most of the cell types other than MGCs and ON-PGC looked squeezed, another UMAP embedding without MGCs were also visualized (bottom right).

**Supplementary Figure 4.**
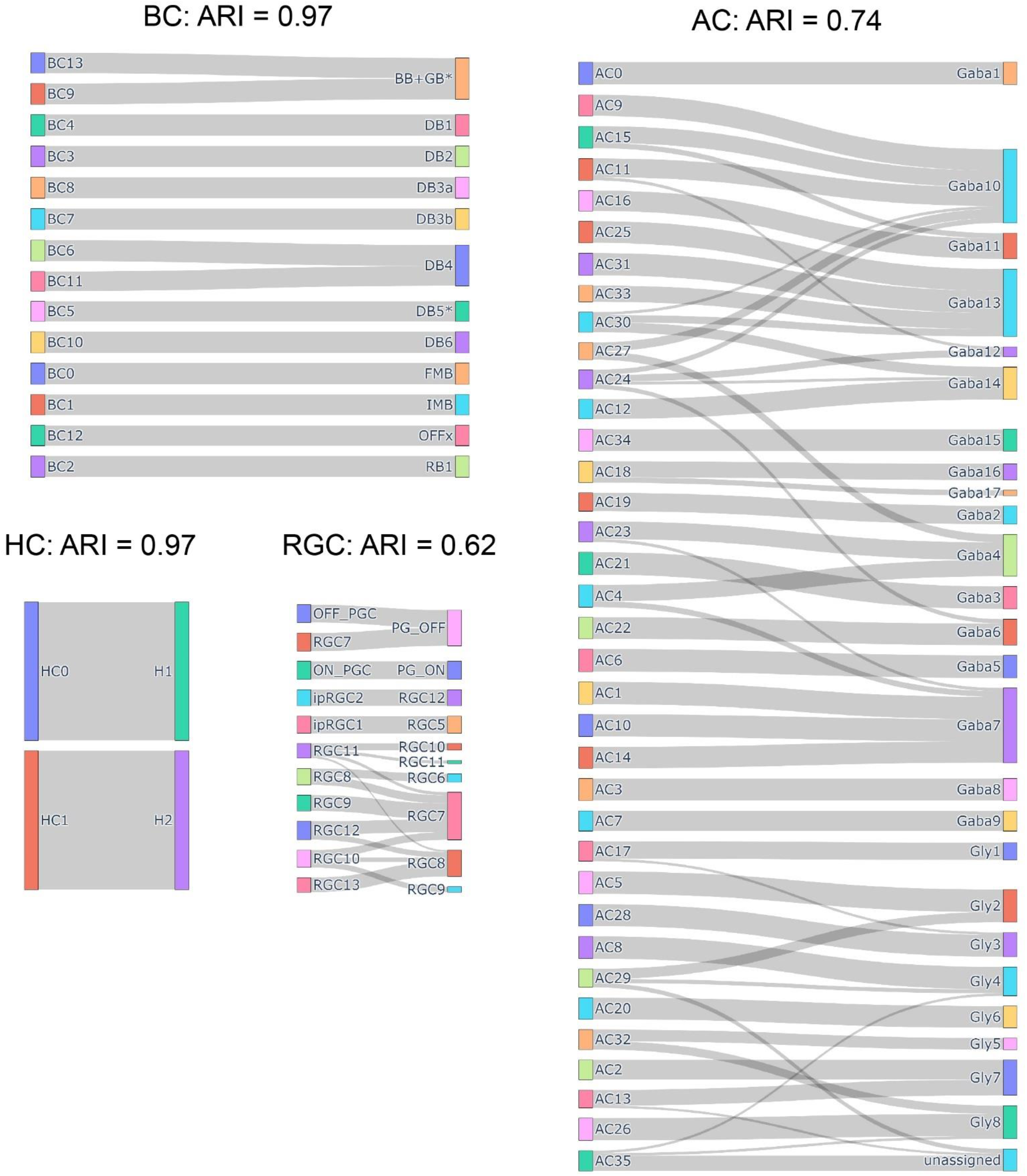
Comparison of the cell type transcriptome profiles between the current study and the previous report by Yan et al. For each cell type, random forest classifier was trained in the Yan dataset and applied to the current dataset for label prediction. The portion of each cluster (in the current work) being predicted as a cell type (in the Yan dataset) were then visualized as Sankey diagrams. Adjusted Rand Index (ARI) were computed between the two label sets (current clustering vs label prediction) to evaluate the consistency of the two datasets.

**Supplementary Figure 5.**
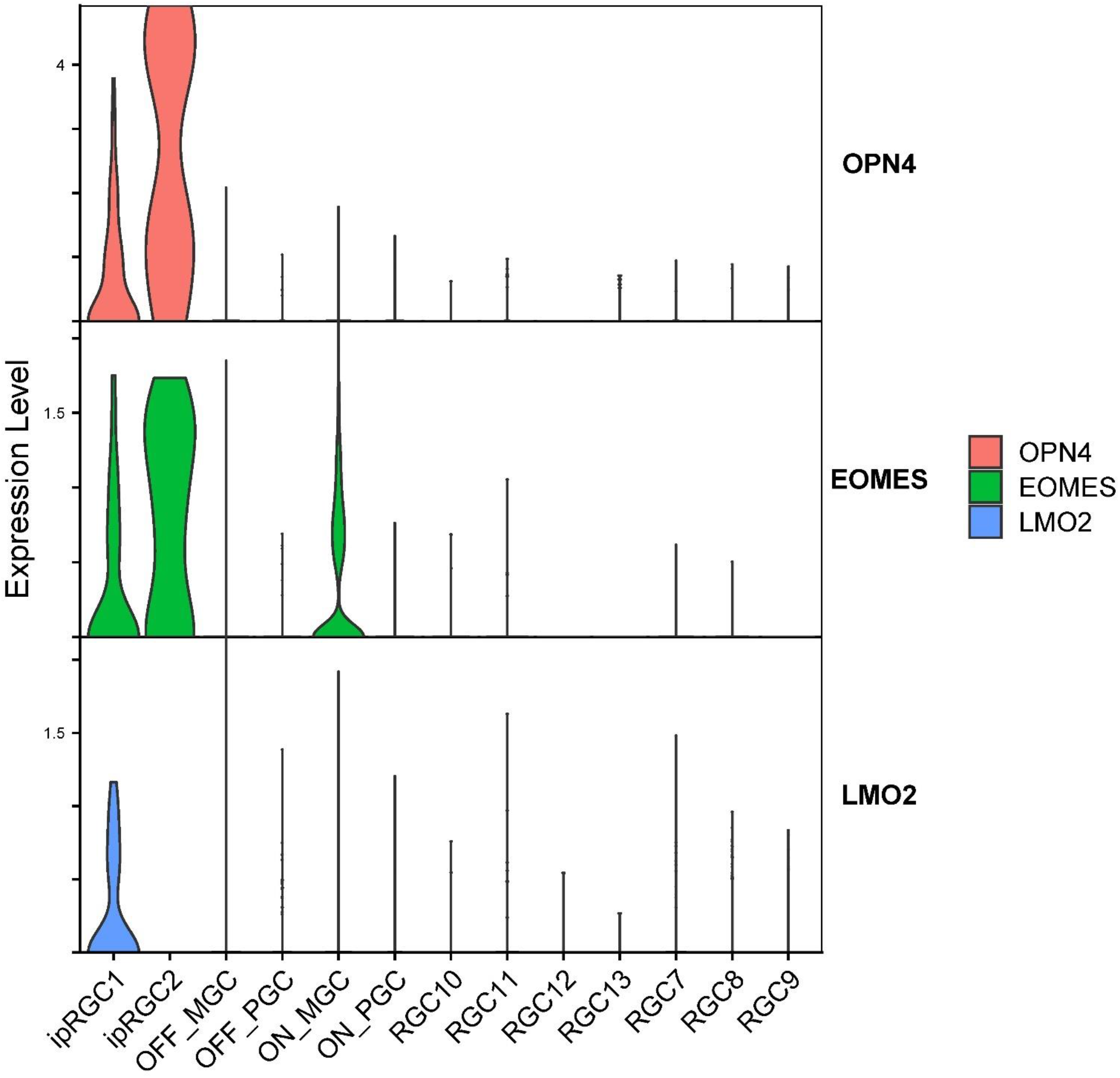
The expression of previously reported OPN4 positive RGC markers in the current dataset.

**Supplementary Figure 6.**
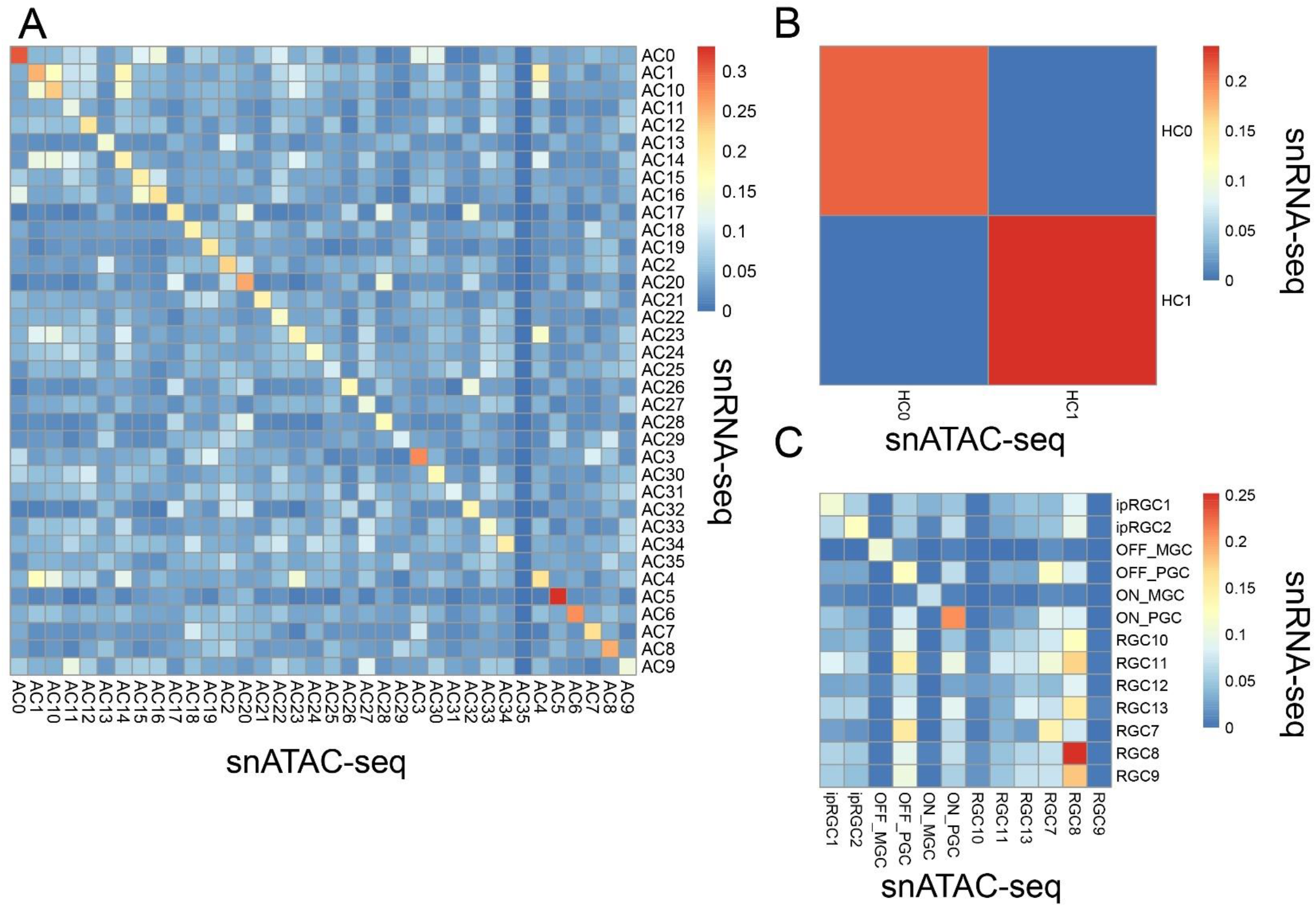
Evaluation of the reliability of the integration of snRNA-seq and snATAC-seq. Heatmap representing the similarity between the differentially expressed genes and the differentially accessible genes from each AC (A), HC (B), and RGC (C) types.

**Supplementary Figure 7.**
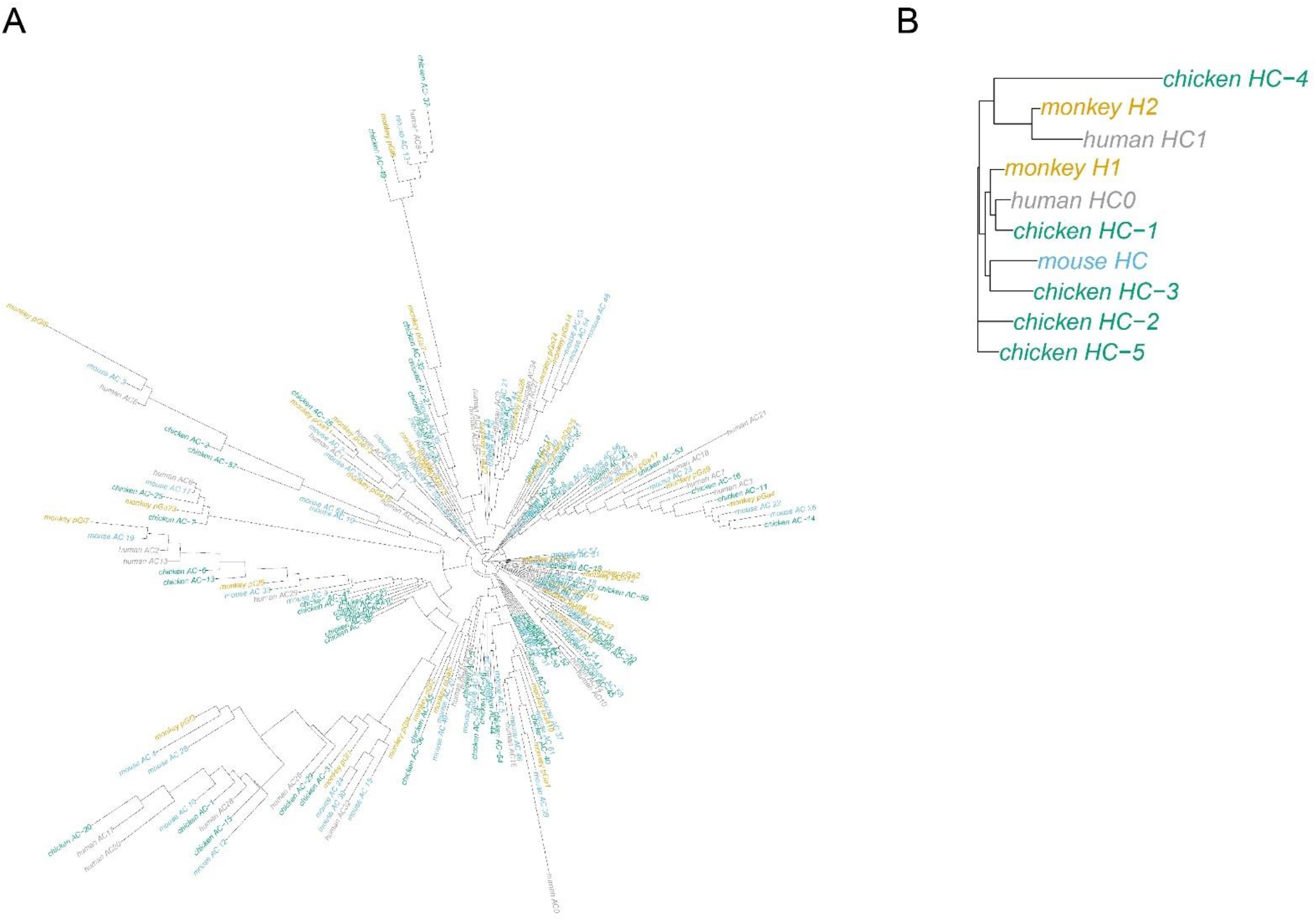
Phylogenetic tree representing the overall similarity of AC (A) and HC (B) types among four species: human (grey), monkey (yellow), mouse (blue), and chicken (green).

**Supplementary Figure 8.**
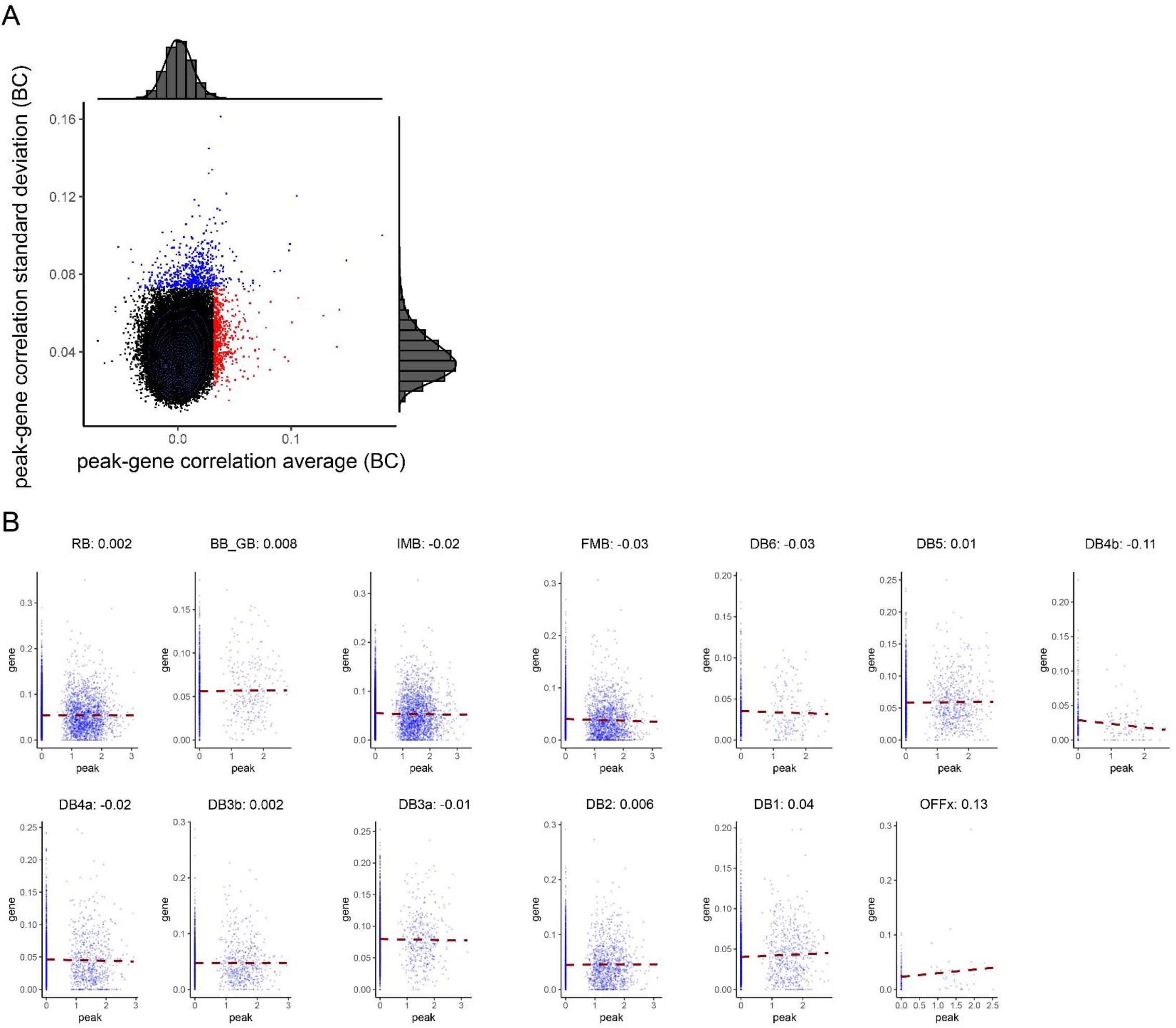
The consistency of CRE functions across types within a major cell class (BC) A. Scatter plot showing the distribution of mean and standard deviation of each peak-gene links among the BC types. The data points being highlighted as blue and red were considered as ‘heterogenous’ and ‘constant’ peak-gene links, respectively. B. The peak gene correlation between chr12:56614952-56617057 and ESYT1 in all bipolar cell types. For each panel, each data point represents a cell and the red dashed-line represents the linear fit of the correlation between the peak and the gene. The Pearson’s correlation coefficients were labeled on each panel.

**Supplementary Figure 9.**
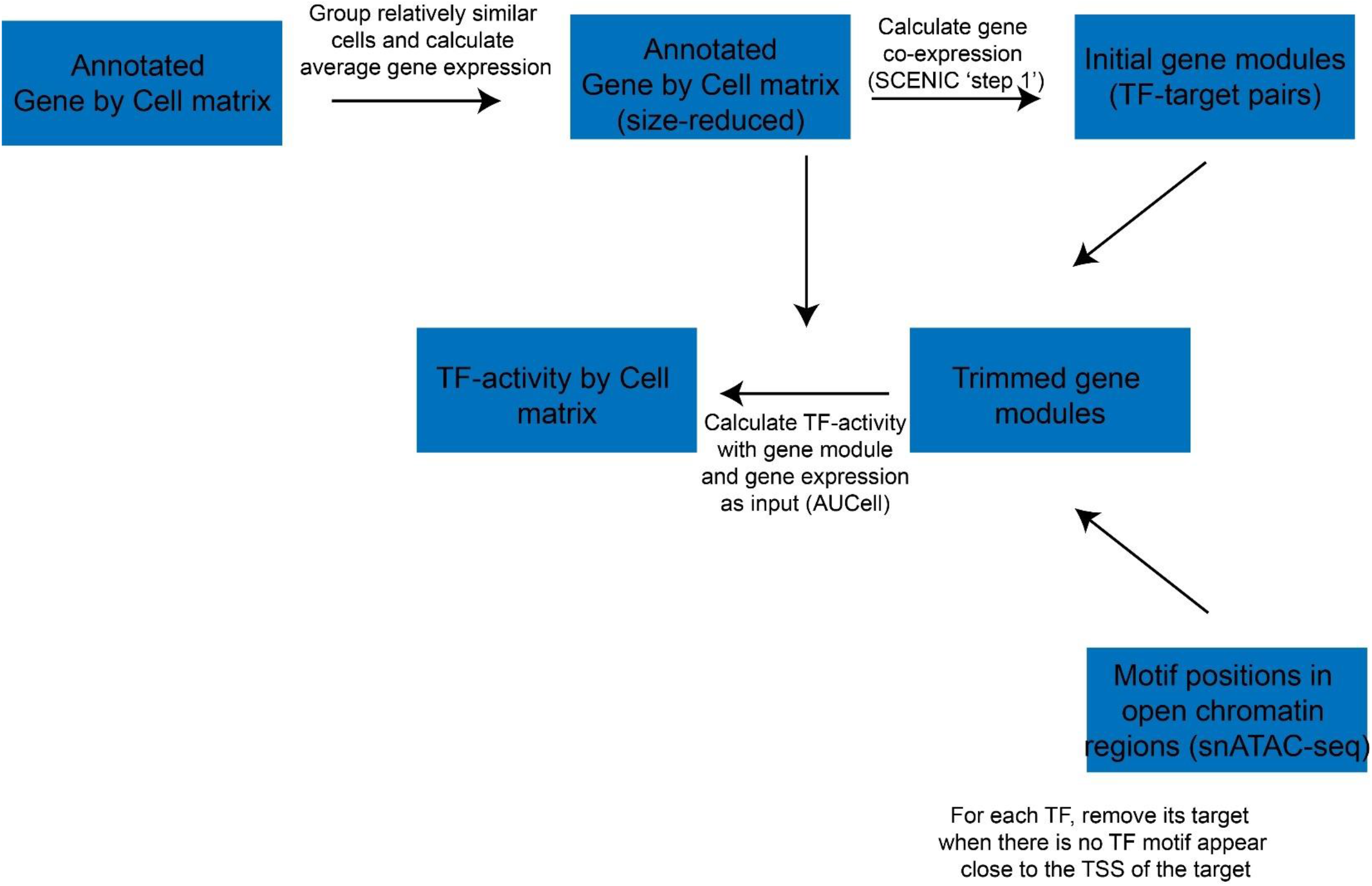
A schematic showing the workflow of the calculation of the TF activity.

**Supplementary Figure 10.**
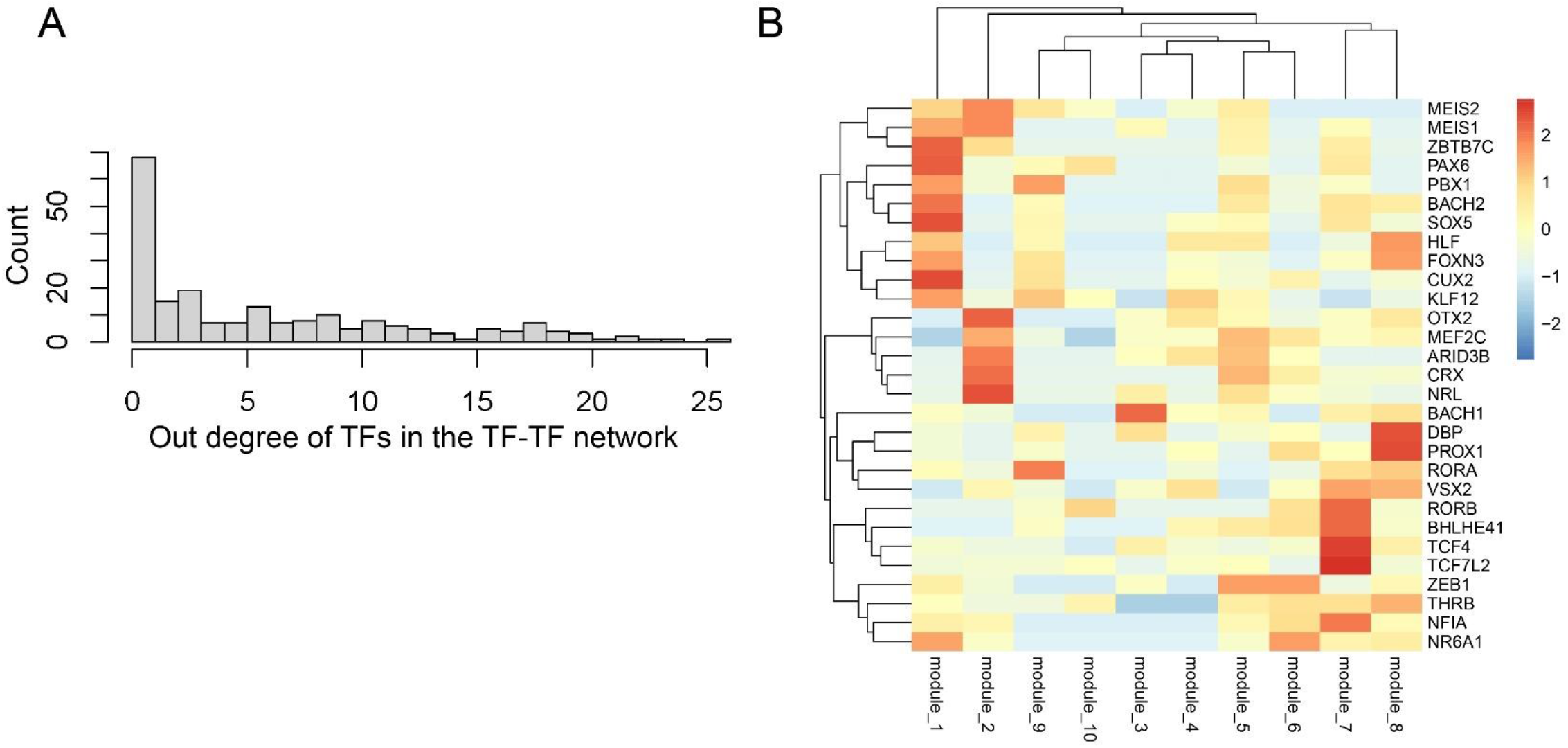
Potential hub regulators of the TF-TF network. A. Histogram showing the distribution of the out degree of TFs in the network. B. Heatmap depicting the overlapping between TF targets and TFs in the pre-defined modules. The TFs were selected based on the out degree (larger than 15). The overlapping was evaluated by computing the Jaccard Similarity. The data was scaled by row before plotting.

